# Enhanced Neuronal Activity and Asynchronous Calcium Transients Revealed in a 3D Organoid Model of Alzheimer’s Disease

**DOI:** 10.1101/2020.07.07.192781

**Authors:** Juan Yin, Antonius M. VanDongen

**Affiliations:** Program in Neuroscience and Behavioural Disorders, Duke-NUS Medical School, 169857 Singapore

## Abstract

Advances in the development of three-dimensional (3D) brain organoids maintained *in vitro* have provided excellent opportunities to study brain development and neurodegenerative disorders, including Alzheimer’s disease (AD). However, there remains a need to generate AD organoids bearing patient-specific genomic backgrounds that can functionally recapitulate key features observed in the AD patient’s brain. To address this need, we successfully generated cerebral organoids from human pluripotent stem cells (hPSCs) derived from a familial AD patient with a mutation in presenilin 2 (PSEN2). An isogenic control hPSC line was generated using CRISPR-Cas9 technology. Both organoids were characterized by analysing their morphology, Aβ42/Aβ40 ratio and functional neuronal network activity. It was found that AD organoids had a higher Aβ42/Aβ40 ratio, asynchronous calcium transients and enhanced neuronal hyperactivity, successfully recapitulating some aspects of AD pathology. Therefore, our study presents a promising organoid-based biosystem for the study of the pathophysiology of AD and a platform for drug development for neurodegenerative disorders.

## Introduction

Alzheimer’s Disease (AD) is the most prevalent cause of dementia, characterized by a progressive loss of both synaptic function and long-term memory formation^1^. Although many efforts have been made to develop therapeutic drugs in the past decades, there is currently no treatment that could prevent, stabilize or reverse the progression of this disease^2^. Therefore, a detailed understanding of the pathophysiology underlying AD and the development of novel therapeutics are highly desirable. However, this remains quite challenging due to the experimental inaccessibility of the functional human brain^3^. Instead, *in vitro* model systems offer alternative and unprecedented opportunities to study AD, as they can recapitulate faithfully some key features involved in AD pathophysiology.

Animal model studies, performed mainly on rodents, have greatly improved our understanding of the mechanisms underlying AD, but these models have flaws as well. Because differences between rodent and human brains are significant, deciphering human diseases using animal models is challenging^4^. For example, mice cannot spontaneously develop an AD phenotype during aging, suggesting that the molecular networks driving AD initiation may be different between humans and mice^4–6^. Neurofibrillary tangles, accelerated phenotypes and sporadic pathology that are observed in human AD brains could also not be recapitulated in animal models^5^. These discrepancies indicate a great demand for paradigms that allow investigations into AD in a “human context”^7^. Advances in human pluripotent stem cells (hPSCs) technology have provided excellent tools to study human neurons derived from AD patients. It is believed that these cells bearing AD patient-specific genetic backgrounds could be used as good models to study AD-associated phenotypes at the molecular and cellular level. For example, several studies have shown that patient-derived hPSCs could produce extracellular amyloid plaques composed of the amyloid beta (Aβ) peptide and intracellular neurofibrillary tangles composed of the microtubule-associated protein tau^8,9^, which are representative of AD pathology. Despite their great potential, two-dimensional (2D) *in vitro* cultures of hPSCs are subjected to some inherent limitations. For example, 2D culture systems fail to reproduce sophisticated cell-cell interactions in the brain^10^. The phenotypes of aberrant extracellular protein aggregation in AD could not be reproduced perfectly, due to the lack of an interstitial compartment in the 2D culture^11^. In addition, cellular diversity, structural complexity and physical architecture could not be reconstructed, because only limited cell types could be generated in 2D culture plates^5^.

To overcome these shortcomings, several groups have developed methodologies to generate cultures of three-dimensional (3D) cerebral organoids. Compared with 2D cultures, organoids are considered to be improved models to study brain development, disease progression and perform drug evaluation, due to their ability to differentiate, self-organize, and form unique, complex, biologically relevant structures^12,13^. The structures formed in organoids are highly reminiscent of the cerebral cortex, which is one of the areas that is severely affected in AD patients. AD organoids generated from familial AD patients with common genetic mutations in presenilin 1 (PSEN1), or the amyloid precursor protein (APP), could successfully recapitulate several key features of AD at the protein and cellular phenotype levels, such as amyloid aggregation, hyperphosphorylated tau protein, endosome abnormalities and neurofibrillary tangles^5,12,13,14^. However, in previously studied AD organoid models, recapitulation of the abnormal functional aspects of AD pathology has been less well studied. Development of such AD organoids models and healthy controls with functional neuronal network activity will be quite useful to study AD pathology. Therefore, we report here our efforts to generate AD organoids bearing a Presenilin 2 (PSEN2) gene mutation from hPSCs derived from familial AD patients and healthy organoids with the same genetic background in which PSEN2 mutation was corrected using CRISPR-Cas9 technology. The presenilin 2 (PSEN2) gene encodes the catalytic components of the γ-secretase enzyme that promotes the final cleavage of APP to generate Aβ^15,16^. The PSEN2 mutation is causally related with the pathogenesis of AD by significantly increasing Aβ42 production, which is prone to intracellular aggregation. So we not only characterized the Aβ42/Aβ40 ratio in the AD organoids to recapitulate this key AD feature, but also studied the effects of PSEN2 mutation on the generation, morphology and function of organoids, offering a new platform to study the mechanism underlying AD pathology and develop drugs.

## Results

### Patient and Control Cells

The linkage of a locus on human chromosome 1q31–42 to early-onset familial Alzheimer’s disease led to the identification of the point mutation PSEN2^N141I^ (422A>T) in the Volga German kindreds in 1995^17^. The AD hPSC lines were collected from a female familial AD patient carrying a heterozygous PSEN2^N141I^ point mutation. The control hPSC lines were collected from a healthy woman who was about the same age as the AD patient. The AD hPSC and control hPSC lines were purchased from Cedars Sinai (ID: CS08iFAD-nxx and CS00iCTR). Both hPSC lines were identified and confirmed by the detection of expression of pluripotency-associated markers, including Nanog and SSEA-1 by immunostaining (**Fig. 1**).

**Figure 1.**
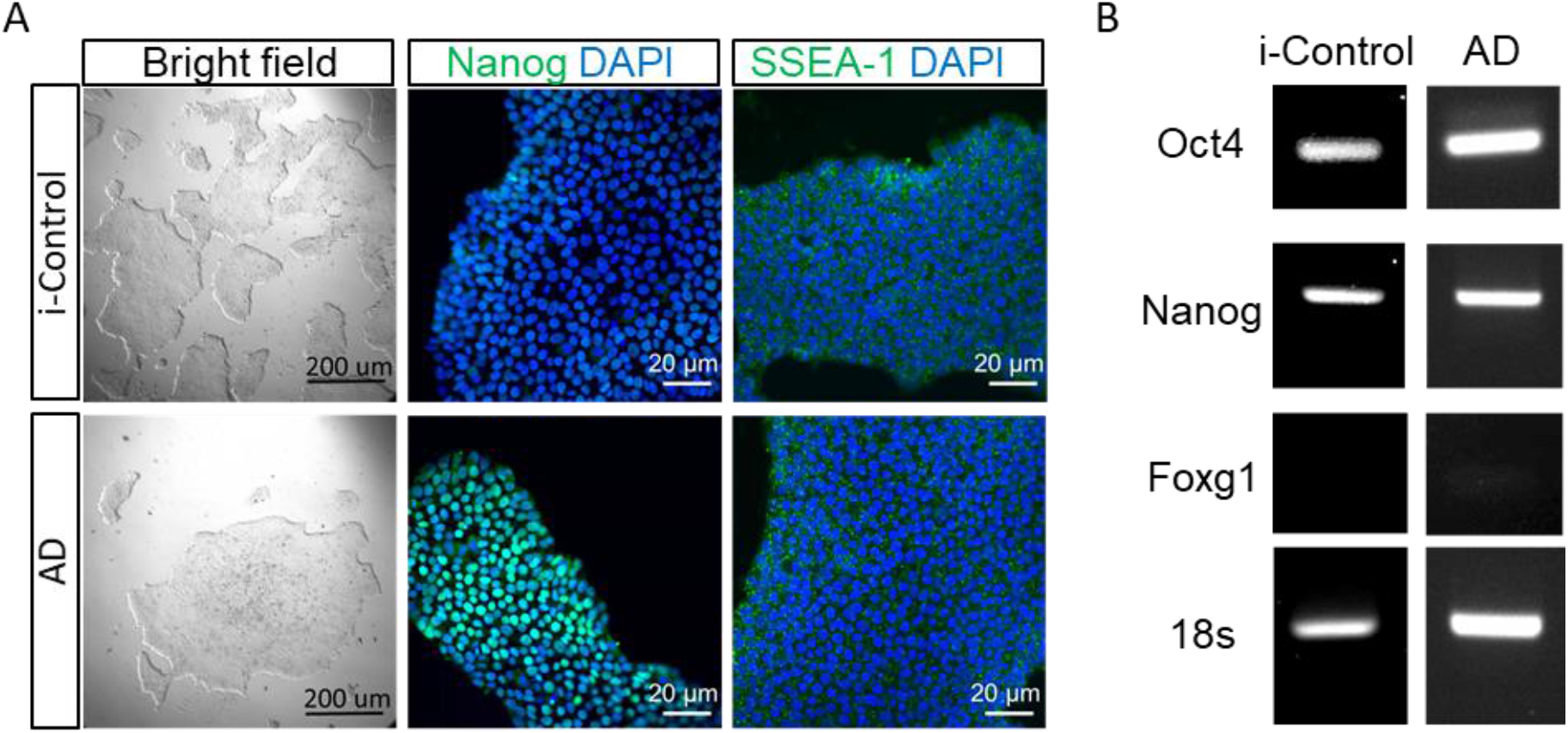
The pluripotent state of hPSCs. (**A**) Bright field images and immunohistochemistry staining of hPSCs. Nucleus was stained with DAPI and hPSCs were stained with Nanog and SSEA-1. (**B**) RT-PRC analysis of hPSC-specific marker (Oct4 and Nanog) and forebrain neuronal marker (Foxg1).

### CRISPR/Cas9-mediated correction of PSEN2^N141I^ mutation in isogenic control cells

To examine the cause-effect relationship between the PSEN2^N141I^ point mutation and AD phenotypes, we generated gene-corrected isogenic control lines (termed as i-Control) from AD hPSC lines using a previously published donor plasmid-mediated CRISPR/Cas9 workflow^18^. In brief, we designed two pairs of sgRNAs that target unique sequences in the introns flanking exon3 of the PSEN2 gene (**Fig. 2A**). The donor plasmid contains the 5’ and 3’ arm, which are the homologous regions of the introns flanking exon3 of the PSEN2 gene, and one PGK-puromycin cassette. We then transfected hPSCs with donor plasmid and the Cas9 nickase (Cas9n) plasmid, along with both sgRNA1 and sgRNA2. After drug selection, PCR genotyping and sequencing showed that about 25% of the clones were targeted to at least one allele (**Fig. 2B, C**). The integrity of the cDNA was verified by RT-PCR and DNA sequencing (**Fig. 2D**). Finally, the successfully corrected clones, whose genomic background was the same as that of the AD hPSC line, were expanded and used to generate organoids (termed as i-Control organoids).

**Figure 2.**
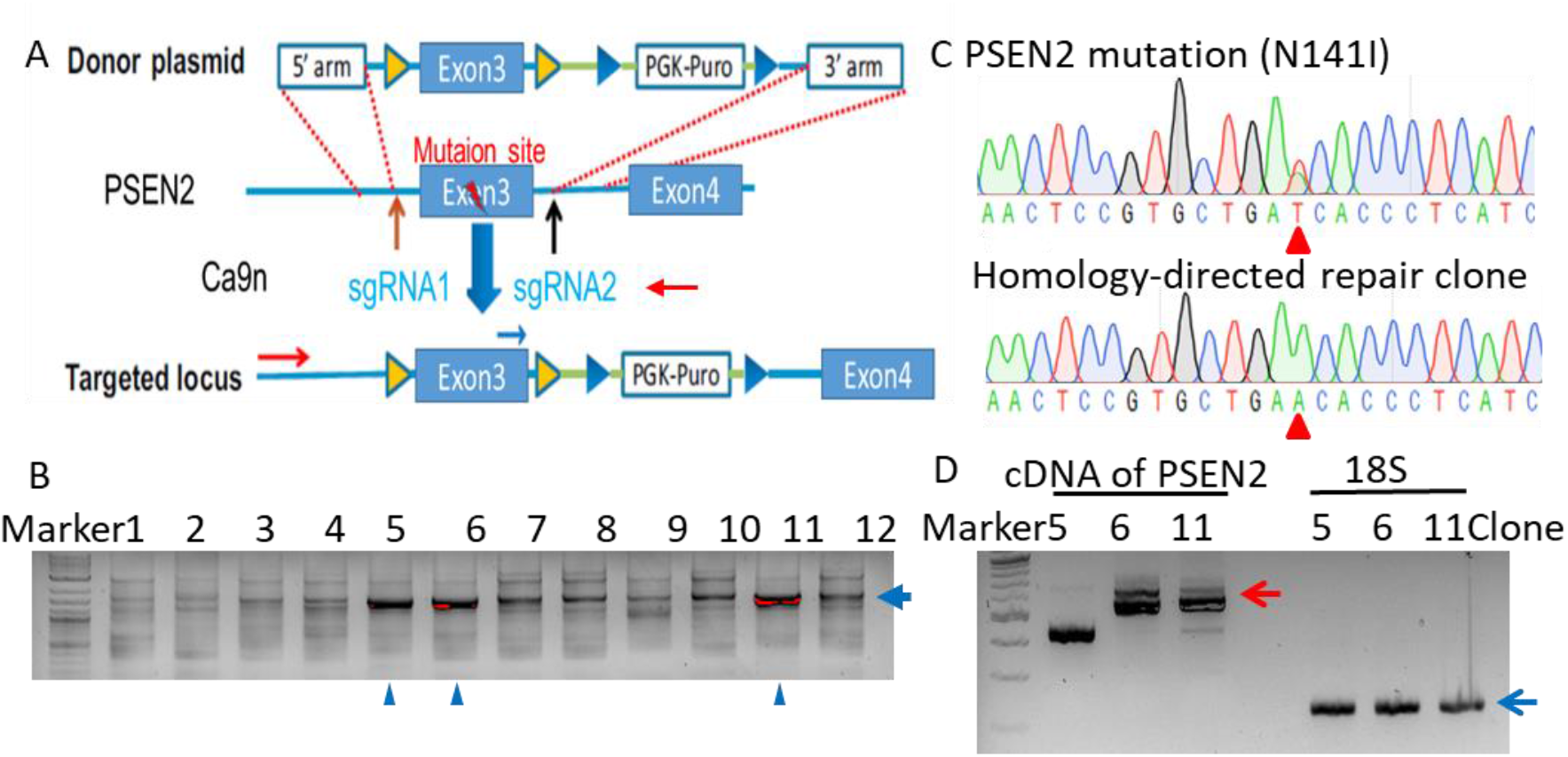
CRISPR/Cas9-mediated correction of PSEN2^N141I^ hPSCs lines. (**A**) Schematic depiction of the targeting strategy for exon 3 of the PSEN2 locus. Vertical arrows indicate sgRNA1 and sgRNA2 targeting sites. The genotyping PCR primer are indicated (red arrows). Donor plasmids: PGK, phosphoglycerate kinase promoter; Puro, puromycin-resistance gene. (**B**) PCR genotyping of hPSCs clones targeted by both sgRNA1 and sgRNA2. PCR products for the correctly targeted PSEN2 locus are 1,300 bp (blue arrows). One targets the intron and the other targets the PGK-Puro. Clone 5,6 and 11 are selected to sequence. (**C**) Representative sequencing results of PSEN2 mutation (upper) and homology-directed repair clone (lower). A heterozygous substitution (A>T at nucleotide 787) in PSEN2 gene results in an Asn141Ile (N141I) missense mutation. (**D**) RT-PRC analysis of cDNA of PSEN2. The red arrow indicated the full length of cDNA. The blue arrow indicated 18s.

### Generation of cerebral organoids

Several protocols have been developed to generate cerebral organoids from hPSCs^19–23^; we followed the protocol which is provided by the Stemcell Technologies. We generated organoids from three hPSC lines, which included two control hPSC lines (the above control line from a healthy individual and the i-Control with the corrected gene mutation, respectively) and one hPSC line bearing the PSEN2^N141I^ mutation. The cerebral organoids were generated from these hPSCs in four stages: embryonic body (EB) formation, neuroectoderm induction, neuroepithelia expansion and organoids maturation. After EB formation, they were fed with induction medium to induce differentiation of the primitive neuroepithelia at 5 days. Then the neuroepithelial tissues were embedded into droplets supported by a 3D matrix composed of Matrigel in which organoids expanded to develop neuroepithelia and showed proliferation of neural progenitor cells. After that, the organoids were transferred to the maturation medium to produce neurons and progenitors in a large continuous cortical tissue within an organoid at 40 days (**Fig. 3**).

**Figure 3.**
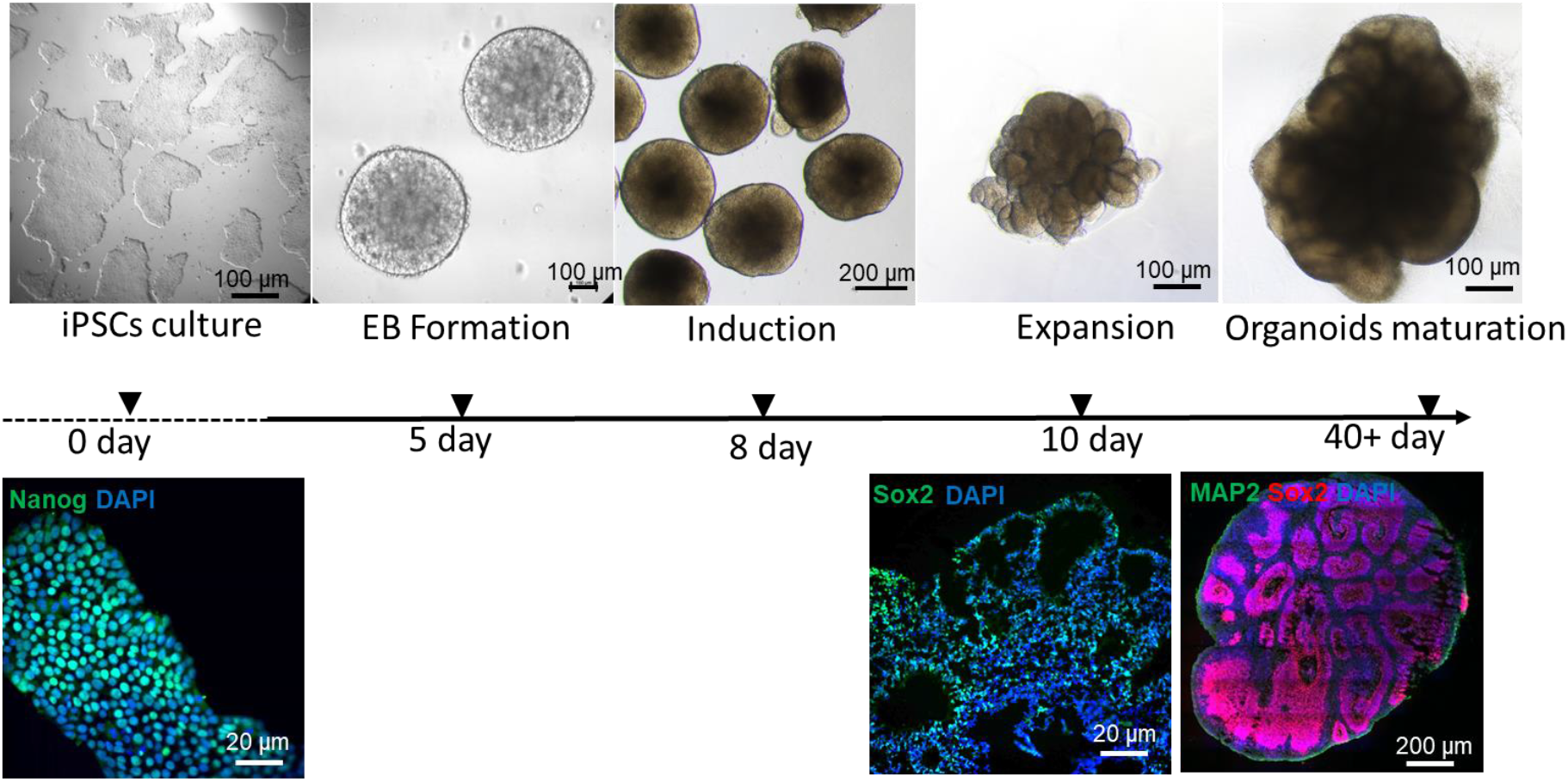
The main stages of the experimental method used to generate human cerebral organoids from hPSCs. Representative immunostaining images (lower panel) showed the pluripotent state of hPSCs (Nanog), neural progenitor cells (Sox2) and neurons (MAP2) in organoids, respectively.

### AD phenotypes in 3D organoids

Immunostaining showed that the organoids generated from AD hPSCs bearing the PSEN2^N141I^ mutation can develop several continual cortical tissues, and their structures are not significantly different from that of organoids generated from control and i-Control hPSCs. At 12 days, almost all the cells in the organoids were neural progenitor cells marked by Sox2 expression. At 25 days, occasional neurons were visible in the organoids, marked by Tuj1 expression. The organoids progressively produced more neurons and thickened cerebral tissues over the subsequent 1 month (**Fig. 4A**).

**Figure 4.**
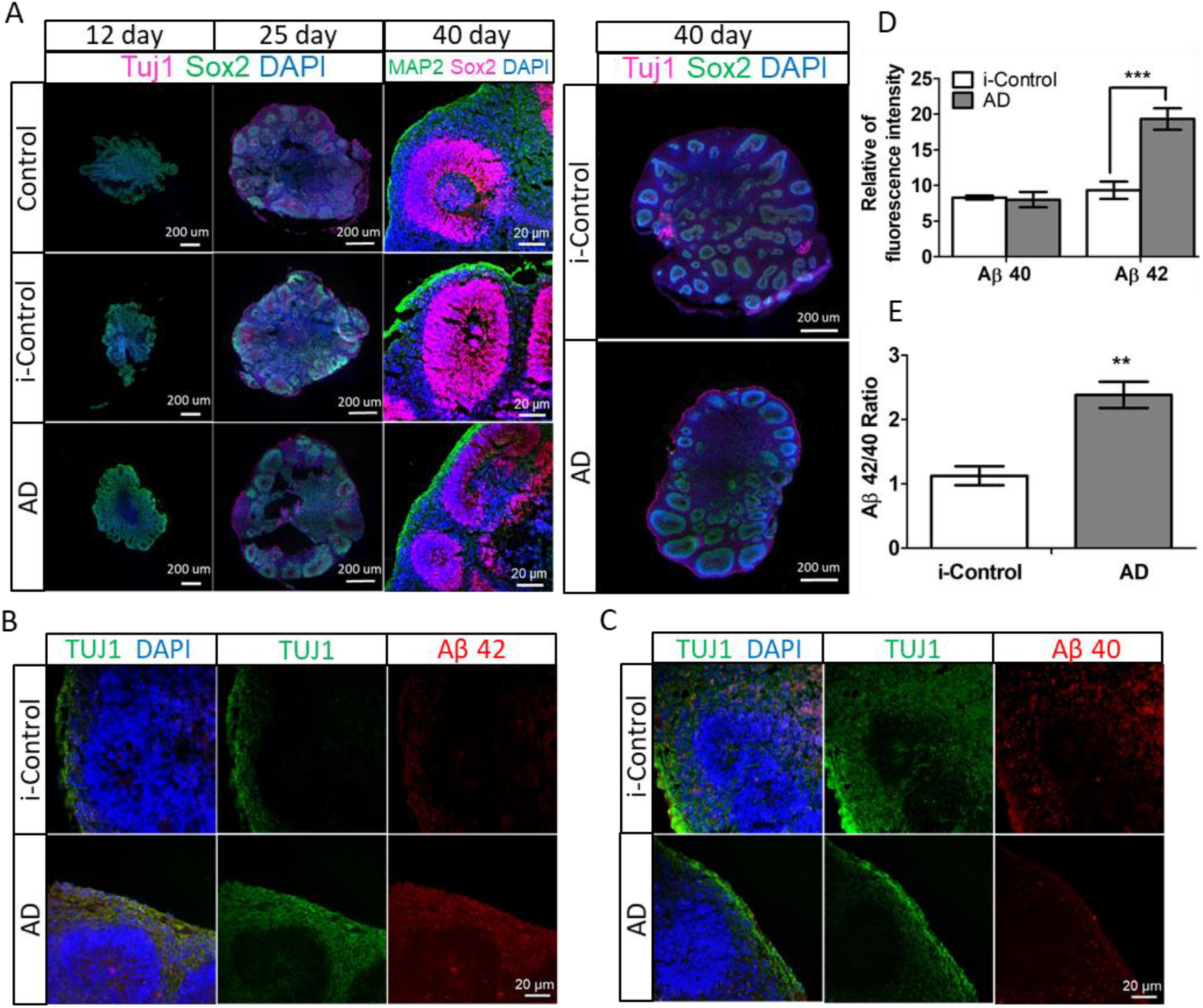
Organoids recapitulate organization of cerebral cortex. (**A**) Sectioning and immunohistochemistry showing complex neuroepithelial tissues containing neural progenitors (SOX2), neurons (TUJ1), and nuclei (DAPI) at day 12, 25, and 40 in the control, i-Control and AD organoids. (**B**) Sectioning and immunohistochemistry showing the expression of Aβ42 at neuroepithelial tissues on day 45. The nucleus was stained with DAPI and neurons were stained with TUJ1. (**C**) Sectioning and immunohistochemistry showing the expression of Aβ40 at neuroepithelial tissues on day 45. Nuclei were stained with DAPI and neurons were stained with TUJ1. (**D**) Quantification of expression level of Aβ40 and Aβ42 (n=3, **P<0.01, ***p < 0.001). (**E**) Quantification of Aβ 42/40 ratio (n=3, **P<0.01).

Previous studies of the effect of the PSEN2^N411I^ mutation in human brain, cerebrospinal fluid, and plasma, as well as in transgenic animals and cellular systems showed that the ratio of Aβ42/Aβ40 was consistently elevated^24^. An increased production of Aβ42 is a major hallmark in the brains of patients with AD, as the higher concentration of Aβ42 oligomers is toxic to nerve cells^16^. Therefore, to determine whether the organoids generated in our work can recapitulate this key pathogenic feature of AD, we tested the expression level of Aβ42, Aβ40 and found that Aβ42 was significantly increased, while Aβ40 was almost unchanged compared to i-Control organoids (**Fig. 4B-D**). Quantification showed that the ratio of Aβ42/Aβ40 was significantly increased (**Fig. 4E**). These data demonstrated that AD organoids bearing the PSEN2^N141I^ mutation could recapitulate an AD-like feature: toxic protein accumulation.

### AD organoids were smaller than i-Control organoids due to an increase in apoptosis

At early differentiation stages, the morphology of organoids generated from control, i-Control and AD hPSCs did not show significant differences. All organoids developed enlarged neuroepithelia, as evidenced by budding at the EB surface. After 40 days, they all exhibited dense cores with regions of organoids displaying optically translucent edges. However, the AD organoids were smaller than organoids generated from control and i-Control hPSCs (**Fig. 5A, B**). Next, to more precisely analyze the effect of the PSEN2 mutation on the defective morphology of AD organoids, i-Control and AD organoids were compared and characterized because they are isogenic. This may minimize the unknown effects caused by different genetic backgrounds of control organoids derived from different healthy individuals.

**Figure 5.**
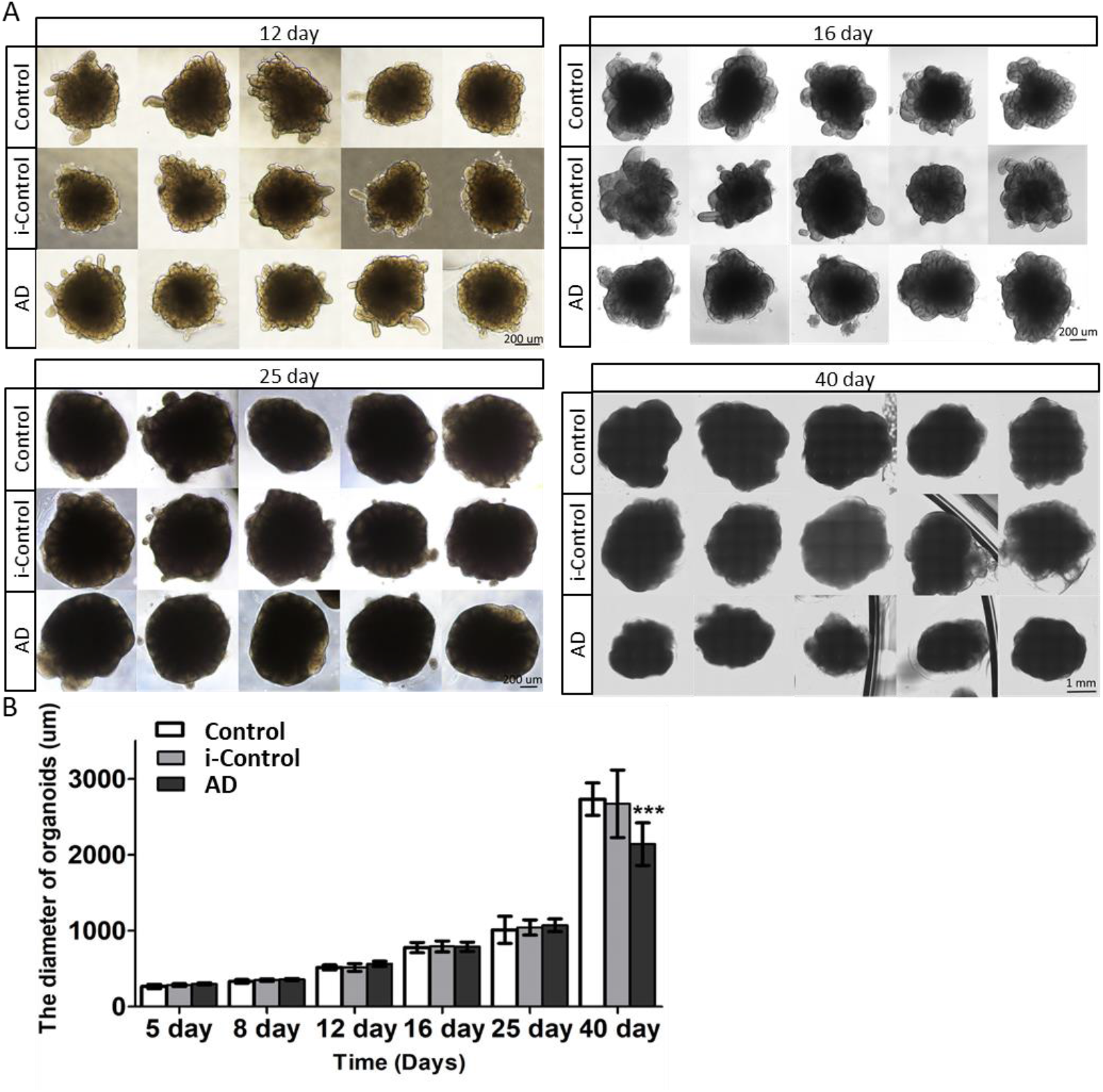
The size of organoids at different development stages. (**A**) Phase-contrast images of organoids at day 12, 16, 25, and 40. (**B**) Quantification of the size of organoids by measuring the diameters of organoids (n=15, ***p < 0.001).

In previous studies, it was found that AD mouse models showed a significant reduction of cell proliferation in neurogenic niches at early stages of development. Therefore, we decided to test whether the smaller size of AD organoids was caused by a reduced cell proliferation. Immunostaining for the expression of cell proliferation markers showed that more cells were positive for the proliferation marker ki67 at an early stage (day 16) than that at later stage (day 45) in both i-Control and AD organoids (**Fig. 6A**). This suggests that there were more neural progenitor cells in the early stages of organoid development as mature neurons lost their ability to proliferate, which is consistent with the result shown in Fig. 3A. But statistical analysis showed that there was no difference in cell proliferation rate between i-Control and AD organoids at the same growth stage (16 or 45 day) (**Fig. 6C**).

**Figure 6.**
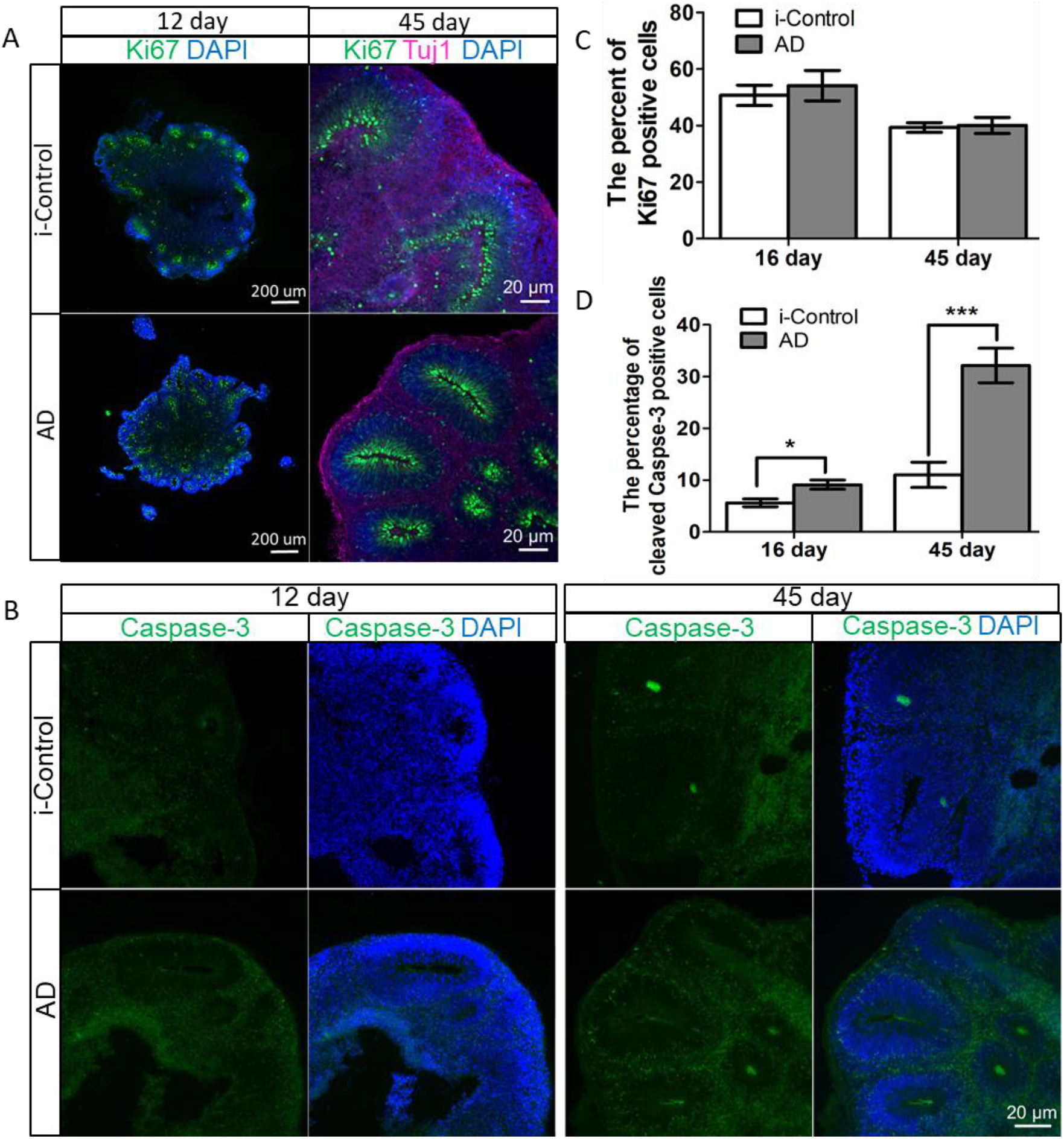
Image analysis of cerebral organoids. (**A**) Sectioning and immunohistochemistry showing the expression of Ki67 at neuroepithelial tissues at day 12 and 45 in i-Control and AD organoids. (**B**) Sectioning and immunohistochemistry showing the expression of Cleaved-Caspase-3 at neuroepithelial tissues at day 12 and 45 in the i-Control and AD organoids. (**C**) Quantification of expression level of Ki67 (n=3) in the i-Control and AD organoids. (**D**) Quantification of expression level of Cleaved-Caspase-3 (n=3, *P<0.05, ***p < 0.01) in the i-Control and AD organoids.

In the clinic, AD is characterized by marked neuronal loss and brain atrophy^25–27^. Previous studies of expression of cell proliferation markers demonstrated that both wild type and mutant PSEN2 could trigger apoptosis in a wide range of cell lines, including human HEK293 cells and murine neurons^28^. To determine whether this phenotype was maintained in AD organoids, we measured the expression of a cell apoptosis marker (cleaved caspase-3) at different growth stages. At 16 days, there only a few cells stained positive for cleaved caspase-3 in both i-Control and AD organoids. But the activity of caspased-3 in the AD organoids was slightly higher than that of in the i-Control. At 40 days, compared with i-Control, neurons exhibited dense caspase-3 staining over a larger area in the AD organoids, showing that the activity of caspase-3 in AD organoids was significantly higher than those derived from i-Control hPSCs (**Fig. 6B, D**). Moreover, the intensity of detected green fluorescence signals inside the region of organoids was much stronger than that in the peripheral region (Fig. 3D), indicating that more healthy neurons were located along the perimeter of organoids while centrally located cells had much higher apoptosis activity, which may be caused by the poor availability of nutrients, oxygen, and growth or neurotropic factor gradients inside the organoids^29^. Therefore, these results show that much higher apoptosis may account for the defect in the morphology of AD organoids, resulting in a smaller size.

### Calcium homeostasis is disrupted in AD organoids

After characterizing their morphology and protein marker expression, we continued to characterize the functional neuronal network activity in the i-Control and AD organoids. Accumulating evidence has shown that PSEN2 mutations can disrupt calcium signalling, resulting in neurodegeneration in AD^30,31^. Therefore, we measured Ca^2+^ oscillations to analyse the calcium homeostasis in both organoids by using a fluorescent calcium indicator (Fluo-4). In order to better observe the cells inside the organoids, we cut the organoid into two halves and plated one half on a culture dish at day 40 (**Fig. 7A**). The neurons migrate and spread out to form a single cell layer around the periphery of the organoids (**Fig. 7B**). Spontaneous calcium dynamics were detected in the migrated neurons after 3 weeks in culture. Imaging data is presented by calculating relative changes in fluorescence (ΔF/F) in regions of interest corresponding to neurons. Interestingly, whereas i-Control organoids displayed synchronized calcium transients typically reported for cultured neuronal networks, the calcium transients from AD organoids lacked synchronization (**Fig. 7C-D**). However, AD organoids were more active than i-Control organoids (**Fig. 7I**). This data suggests that the PSEN2^N141I^ mutation may enhance neuronal hyperactivity. At the same time, we found that the amplitudes of spontaneous Ca^2+^ transients organoids containing the in PSEN2^N141I^ mutation were significantly higher than those in the i-Control organoids (**Fig. 7H**). To further confirm that this functional behaviour was caused by the organized 3D cell assembly, the floating organoids were re-dissociated into single cells to generate a monolayer culture with the purpose to test the Ca^2+^ transients at the 2D cell level (**Fig. 7E**). However, after 3 weeks of culture, the calcium homeostasis patterns displayed in the organoids were lost in 2D. There were no significant differences in the activity patterns, such as the synchronicity and numbers of spikes, between i-Control and AD neurons (**Fig. 7F, G**). Moreover, the frequency of Ca^2+^ spikes was also significantly lower in the 2D cultures than in the 3D organoids. Taken together, these data indicate that neuronal networks are better organized in organoids than in monolayer cultures, as more spontaneous Ca^2+^ transients could be detected in organoids. This experimental evidence indicates that organoids have great potential to be used as a functional and “information-rich” *in vitro* model to study AD.

**Figure 7.**
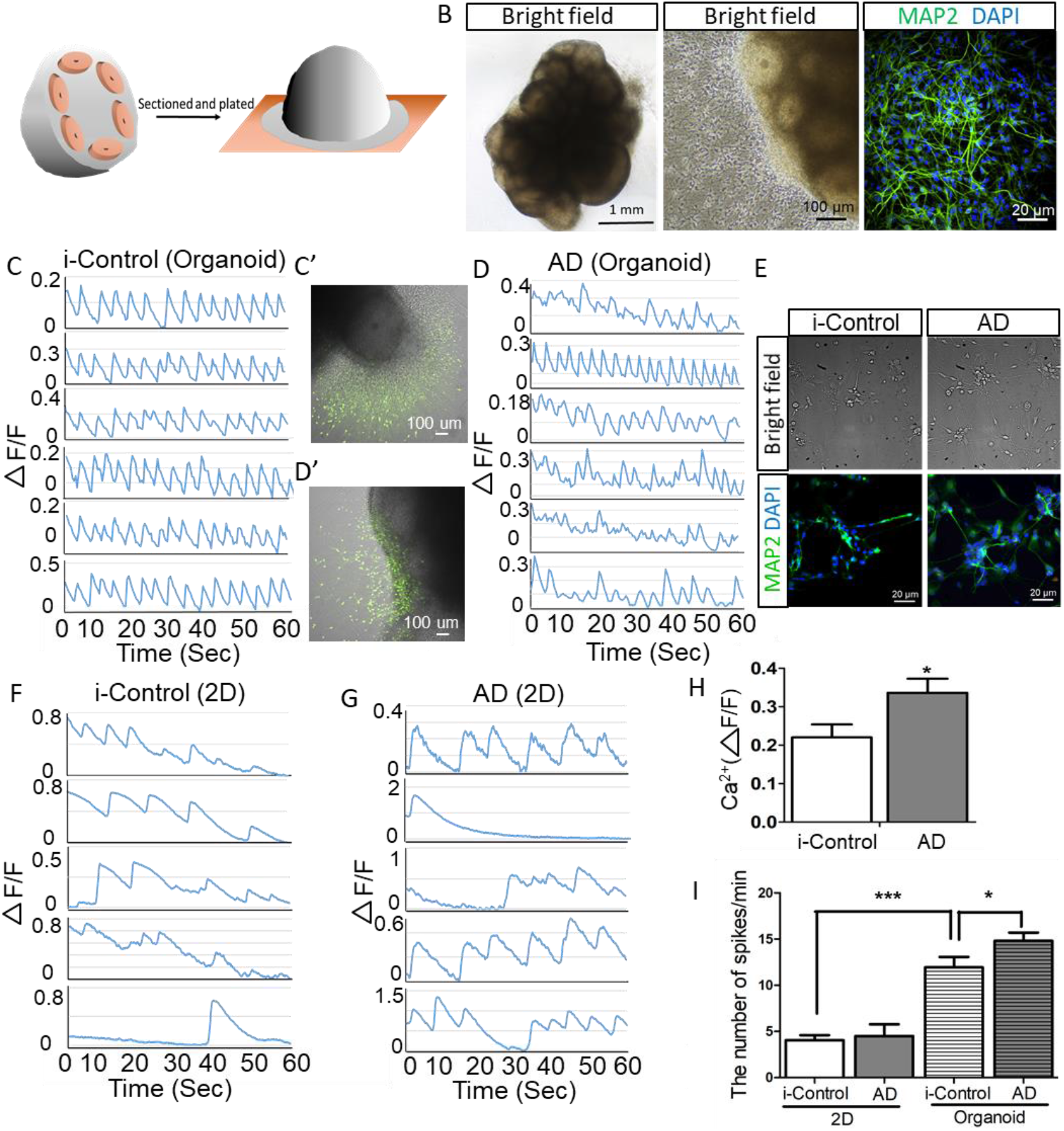
Analysis of calcium activity in cerebral organoids. (**A**) Schematic depiction of cutting the organoids and plating on half of it on the plate at day 40. (**B**) Left panel: Bright field images of whole organoids. Middle panel: Bright field image of half of an organoid that was cultured on a 35 mm glass bottom dish for 3 weeks showing neurons that have migrated out from the surface of the organoid. Right panel: immunohistochemistry staining of neurons migrating away from the organoids. Nuclei were stained with DAPI and neurons were stained with TUJ1. (**C**) Single-cell tracings of spontaneous calcium surges in the migrated neurons from isogenic control organoids (regions of interest shown in C’) as measured by change in fluorescence (arbitrary units). (**D**) Single-cell tracings of spontaneous calcium surges in the migrated neurons from AD organoids (regions of interest showed in D’) as measured by change in fluorescence (arbitrary units). (**E**) Bright field images and immunohistochemistry staining of neuronal cells which were dissociated from organoids. Nuclei were stained with DAPI and neuronal cells were stained with MAP2. (**F**) Single-cell tracings of spontaneous calcium surges in dissociated neurons from isogenic control organoids. (**G**) Single-cell tracings of spontaneous calcium surges in dissociated neurons from AD organoids. (**H**) Quantification of average normalized calcium responses for baseline in isogenic control organoids and AD organoids (n=5, *P<0.05). (**I**) Quantification of the numbers of spikes per minute (n=15, *P<0.05, ***p < 0.01).

### Effect of drug treatment on spontaneous Ca^2+^ transients in organoids

To further analyse the effect of the PSEN2^N141I^ point mutation on dynamic changes of the spontaneous Ca^2+^ transients, two types of drugs were used (**Fig. 8A**). 4-AP is an A-type potassium channel blocker that can induce epileptiform activity in the rat hippocampus^32^. In our assay, we observed that the frequency of spikes was increased about 2-fold by addition of 10 μM 4-AP in both i-Control and AD organoids, but there was no significant difference between them before or after drug treatment (**Fig. 8B**). Bicuculline methochloride is a GABA receptor antagonist which increases the activity of neuronal networks in organoids^33^. Similar to the organoids treated by 4-AP, the frequency of spikes was also increased about 2-fold after adding 50 μM Bicuculline methochloride, but there was also no obvious difference in i-Control and AD organoids before or after drug treatment (**Fig. 8C**). These results suggest that although both 4-AP and Bicuculline methochloride can induce the spontaneous Ca^2+^ transients in both organoids, PSEN2 mutations may have no significant effects on their drug sensitivity.

**Figure 8.**
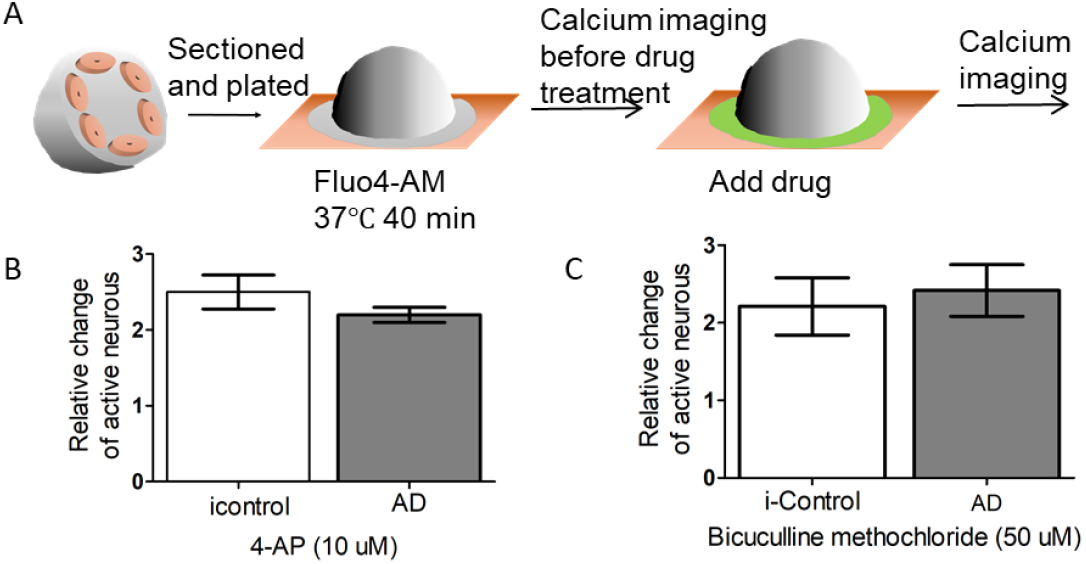
Analysis of Calcium Activity in organoids after drug treatment. (**A**) Schematic depiction of the drug load test. (**B**) The relative number of activate neurons after 4-AP treatment compared to that before drug treatment (n=15). (**C**) The relative number of activate neurons after Bicuculline methochloride treatment compared to that before drug treatment (n=15).

## Discussion

In this contribution, we have successfully generated cerebral organoids from AD patients-derived hPSCs bearing the PSEN2^N141I^ point mutation and an isogenic control in which the mutation was reversed by using the CRISPR/Cas9 system, providing evidence that organoids with the PSEN2^N141I^ point mutation could recapitulate several key features of AD pathology, such as an increased Aβ42/Aβ40 ratio and dysfunctional neuronal network activity.

Previous studies on organoid modelling of brain development and neurodegenerative disorders revealed that certain types of disease-derived organoids exhibited a smaller overall organoid size compared to control organoids^34^. For example, organoids derived from Miller-Dieker syndrome, autosomal recessive primary microcephaly, and Aicardi-Goutieres syndrome were smaller than the control organoids, which was caused by a decrease in neural progenitor proliferation or an increase in neuronal apoptosis^35–37^. Unexpectedly, in our study, we found that a morphological defect occurred in AD organoids after 40 days in culture. Compared to i-Control organoids, AD organoids were smaller in size, which was shown to be caused by an increase in neuronal cell death. We also detected a higher expression of caspase-3 in the AD organoids, indicating an enhanced level of apoptosis. Thus, this disease model may be useful to study the potential mechanism of caspase-3 in progressive synaptic degeneration and neuronal loss in AD in future studies.

Normal calcium signalling is important to maintain brain development and function^38–40^. Destabilized calcium signalling seems to be central to the pathogenesis of Alzheimer’s disease, and targeting this process might be therapeutically beneficial^31^. To investigate this process, generation of organoids that can recapitulate the functional neuronal network activity in human brain has attracted considerable interests. Although it is challenging, several recent studies have reported inspiring results. For example, one study showed that the cerebral organoids not only mimicked early human neurodevelopment at the cellular and molecular level, but they also form neuronal networks which could display periodic and regular oscillatory events^41^. However, experimental recapitulation of the disruption of calcium homeostasis in AD organoids remains still underappreciated. To fill this gap in knowledge, we have analysed neuronal network activity in AD organoids and isogenic controls, respectively. We found that synchronized Ca^2+^ transients could be observed in the neurons that migrated out from control organoids in which the PSEN2^N141I^ mutation was corrected by CRISPR/Cas9 technology. This result was consistent with a previous study, which showed that cerebral organoids derived from a healthy individual produced self-organized neuronal networks and exhibited spontaneous Ca^2+^ transients in synchronized patterns^42^. More importantly, in the current work we found non-synchronized Ca^2+^ transients in the neurons that had migrated out from the AD organoids carrying the PSEN2^N141I^ mutation, which was an obvious contrast compared to the control. This data recapitulates findings from clinical studies that showed diminished fluctuations in the level of synchronization in neural circuits from AD patients^43,44^. Therefore, our data reveal a possible role that the PSEN2^N141I^ mutation may play in the abnormal regulation of calcium homeostasis in AD. Previous studies on cultured hippocampal networks demonstrated that high levels of Aβ42 significantly inhibited the synchronized spontaneous cytoplasmic Ca^2+^ transients^45^. In our study, we also found that the expression level of Aβ42 was significantly increased in AD organoids, which may also have contributed to the loss of synchronization of the Ca^2+^ transients. This aspect needs to be further investigated. Hideya Sakaguchi and his colleagues reported the formation of self-organized and synchronized neuronal network following a dissociation culture of cerebral organoids^42^. However, in our work, once organoids were dissociated into single cells, the frequency of calcium spikes was significantly reduced and synchronous calcium transients disappeared, indicating that the neural activity in 3D organoids may be more mature than that in 2D monolayer cultures.

Neuronal hyperactivity emerges early in the pathological progression of AD, in both mouse models and patients^46^. Clinical studies showed that seizures are more frequent in patients with AD compared to healthy individuals and that they can hasten cognitive decline^47^. To study the potential underlying mechanism, a mouse model of AD using *in vivo* Ca^2+^ imaging revealed that AD-mediated enhancement of neuronal hyperactivity occurred mainly due to a Presenilin-mediated dysfunction of intracellular Ca^2+^ stores^48^, while activity patterns of cortical neurons revealed that AD-related mutations augment neuronal hyperactivity^49^. But whether this is true also in *in vitro* models remains unknown. In this paper we have used cerebral organoids as a model to study neuronal activity by using Ca^2+^ imaging. Our results show that AD organoids carrying the PSEN2^N141I^ mutation displayed enhanced neuronal activity compared to the isogenic control. Therefore, consistent with previous studies on patients and rodent models, this work offers an alternative to study the relationship between Presenilin and neuronal hyperactivity, which can avoid surgical procedures and ethical disputes involved in *in vivo* imaging on living animals.

Taken together, our work has advanced our understanding of functional neuronal network activity in AD bearing a PSEN2 mutation and provided a new organoids-based *in vitro* biosystem to model human AD disease at different levels. We believe that this model will find wide applications, such as the study of mechanisms underlying the pathophysiology of AD and the development of therapeutic drugs for this disease.

## Methods

### Maintenance and characterization of hPSCs

hPSCs were obtained from the Coriell Institute for Medical Research, under their consent and privacy guidelines as described on their website (http://catalog.coriell.org/). The hPSCs carrying PSEN2^N141I^ point mutation is CS08iFAD-nxx (https://web.expasy.org/cellosaurus/CVCL_YX93) and control hPSCs is CS00iCTR. iPSC colonies were grown on Matrigel (Catalog #354277, Corning) in TeSR medium (mTeSR™1, Catalog # 85850, STEMCELL Technologies) according to the manual. Pluripotency was assessed by immunostaining with surface and nuclear pluripotency markers (SSEA-4, Oct4 and Nanog, Catalog #ab109884, Abcam) for subsequent imaging.

### CRISPR/Cas9-mediated correction of the PSEN2^N141I^ mutation

The gene-corrected isogenic control lines (i-Control) were generated from AD hPSC lines using a previously published donor plasmid-mediated CRISPR/Cas9 protocol^18^. The sgRNA vector (Cas9 sgRNA vector, Plasmid #68463), Donor plasmid (PL552, Plasmid #68407), and Cas9 nickase plasmid (pCas9D10A_GFP (Plasmid #44720) were purchased from Addgene. We designed two pairs of sgRNA, which target the intron beside the exon3. The targeting site of sgRNA pair 1A is: ATGCAAAAATTAGCCAGGTG. The targeting site of sgRNA pair 1B is: AGGGTCTCGCCACATTTCCC. The targeting site of sgRNA pair 2A is: CAGGAGTTCAAGTCTAGCCT. The targeting site of sgRNA pair 2B is: TCCCAAGTAGTTGGGACCAC. The donor plasmid, two pairs of sgRNA vectors and Cas9 nickase plasmid were transduced into the AD hPSCs by using Human Stem Cell Nucleofector^®^ Kit 1 (Lonza VPH-5012). Three days after electroporation, puromycin (0.5 μg/ml, Invivogen) was added into the AD hPSCs medium for selection for two weeks. Then individual colonies were picked up and identified by genomic PCR.

### Genomic DNA extraction and Genomic PCR

Genomic DNA was extracted with QuickExtract™ DNA Extraction Solution (Epicentre, QE09050). Genomic PCR was carried out using Q5^®^ High-Fidelity DNA Polymerase (New England Biolabs, M0492L).

### Generation of organoids

The cerebral organoids were generated from hPSCs using STEMdiff™ Cerebral Organoid Kit (Catalog #08570, STEMCELL Technologies) according to the protocol provided by STEMCELL Technologies, and only a small part of the method is modified. In briefly, hPSCs were dissociated to single cells in Dispase (1 U/mL, Catalog #07923, STEMCELL Technologies), and quickly reaggregated using low-cell-adhesion-coated 96-well plates with V-bottomed conical wells (Corning) in EB Formation Medium (12,000 cells per well, 100 mL) containing 10 μM Y-27632. Fresh EB Formation Medium without Y-27632 was changed every other day. At 6 days, EBs were moved to 24-well ultra-low attachment plates (Corning) in Induction Medium. At 8 days, EBs were embedded with Matrigel (Catalog #354277, Corning) and changed to 6-well ultra-low attachment plates (Corning) in Expansion Medium. At 11 days, the medium was replaced with Maturation Medium and plates of organoids were placed on an orbital shaker in incubator (37°C, 5% CO2).

### Dissociation of organoids into single cells

Whole cerebral organoids were washed three times in 1× PBS and dissociated in 2 mL of Accutase (StemPro) containing 0.2 U/μL Dnase I (Roche) for 45 min at 37°C. Cells were collected by centrifugation at 300 × g for 5 min and resuspended in 10 ml of Neural culture media (Neurobasal medium, 1× B-27, 200 mM L-glutamine, 50 U/ml penicillin and 50 mg/ml streptomycin). Then centrifugation was performed at 300 × g for 5 min, and cells were resuspended in Neural culture media. Cell viability was assessed by Trypan blue staining (typically 85– 95% viable) and counted using an automatic cell counter (Bio-Rad). Cells are platedon 100 mg/mL poly-L-ornithine and 10 mg/ mL laminin coated plates at 50,000 cells/cm2 and incubated overnight. The medium was changed twice a week.

### Cryosectioning and immunostaining for organoids

Organoids were fixed with 4% paraformaldehyde overnight at 4 °C and then transferred to 30% sucrose solution. After organoids sank in the sucrose solution overnight at 4 °C, they were embedded in O.C.T. (Sakura, Tokyo, Japan) and sliced in a cryostat (20 μm slices). Following air drying for 30 min, the slides containing the sliced samples were permeabilized with 0.1% triton X-100 for 20 min, and blocked with 10% donkey serum in PBS for 1 hour at room temperature. The slides were incubated with Sox2 antibody (Catalog # AB5603, Chemicon, 1:300), anti-Tuj1 antibody (Catalog #ab7751, Abcam, 1:1000), Ki67 antibody (Catalog #66155, Abcam, 1:300), Cleaved Caspase-3 (Asp175) Antibody (Catalog #9661T, Cell Signaling Technology) for 1 hour at room temperature. Next, the slices were washed with PBST (PBS with 0.05% triton X-100) and incubated with anti-mouse antibody (Thermofisher, Rabbit anti-Mouse IgG (H+L) Secondary Antibody, Alexa Fluor 647, 1:300) or anti-rabbit antibody (Thermofisher, Goat anti-Rabbit IgG (H+L) Secondary Antibody, Alexa Fluor 488, 1:300) for 30 min at room temperature. The nuclei were stained using the DAPI solution (1 μg/mL). The slides were mounted and analyzed using a fluorescence microscope.

### Immunohistochemistry

For immunostaining of hPSCs and neurons, cells were fixed with 4% PFA directly on the 35 mm cell culture dish for 1 hour, and then washed three times with PBS. The cells were treated with PBS containing 1% Triton X-100 for 15 min. Then, cells were incubated in the blocking solution (PBS with 0.1%Triton X-100 plus 10% Donkey serum) for 0.5 h at room temperature. The anti-Oct4 antibodies (Catalog #ab109884, Abcam, 1:20), anti-Nanog antibody (Catalog #ab109884, Abcam, 1:20), anti-SSEA4 antibody (Catalog #ab109884, Abcam, 1:20) or anti-Tuj1 antibody (Catalog #ab7751, Abcam, 1:1000) was diluted in the blocking solution, which was then added to the culture dish for incubation for 1 h at room temperature. Cells were washed four times with PBST (PBS with 0.1% Triton X-100) and then treated with an anti-mouse antibody (Thermofisher, Rabbit anti-Mouse IgG (H+L) Secondary Antibody, Alexa Fluor 488) or anti-rabbit antibody (Thermofisher, Goat anti-Rabbit IgG (H+L) Secondary Antibody, Alexa Fluor 488) diluted at 1:300 in blocking solution for 1 h at room temperature. Then cells were washed four times with PBST and incubated with DAPI (1 μg/mL, diluted in PBS) for 10 min at room temperature for nuclear counterstain. Cells were visualized using a confocal microscope under 20 x magnification.

### Calcium Imaging of the Neuronal Network

For calcium dye loading, the cells were incubated with 10 mM Fluo-4-AM solution (Catalog #F14201, Invitrogen) and 0.04% Pluronic F-127 (Catalog #P3000MP, Invitrogen) for 1 hour in an incubator (37°C, 5% CO2). Excess dye was removed by washing with culture medium three times. Imaging was carried out at 37°C and 5% CO2 using a confocal microscope (Nikon). Time-lapse image sequences were acquired at no intervals for 2 min. The fluorescence change over time is defined as ΔF/F = (F/F_basal_)/F_basal_, where F is the fluorescence at any time point and F_basal_ is the minimum fluorescence of each cell. A neuron was considered active if calcium transients were observed at least once during total imaging period. For pharmacological experiments, bicuculline methochloride (50 mM) and 4-AP (10 mM) was added after 2 min of time-lapse image, imaging data were taken another 5 min after drug treatment in the same field.

### Statistical Analysis

All statistical analyses were performed using the unpaired t-test. Data in graphs are expressed as mean values ±S.E.M. Error bars represent S.E.M.

## Acknowledgments

This work was supported by grant NMRC/OFIRG/0019/2016 from the National Medical Research Council in Singapore.

